# Quantitative analysis and medium components optimizing for culturing a fastidious bacterium *Christensenella minuta*

**DOI:** 10.1101/632836

**Authors:** Hongshi Xiao, Binghuan Liu, Jie Yong, Haiyan Zhou

## Abstract

*Christensenella minuta* is a heritable bacterium with controversial physiologies associating with both obesity and potential pathogenicity. Since this bacterium is fastidious to culture, it is hardly to well understand its biological feature. We develop a strategy for statistical analysis of this low abundant strain and optimize culture condition to make a significant improvement on its biomass and facilitate the researches about the metabolism and function of this bacterium. Basing on the fluorogenic quantitative technology, a quantitative approach was successfully constructed for *Christensenella minuta* by plotting *C*t value from fluorescence quantitative PCR against the logarithm of concentration gradient of plasmids containing 16S rDNA of the strain. This method exhibited to have specificity on analyzing the strain biomass statistically. For improving the strain biomass, “komodo” predicted to optimize medium components and metabonomics analysis explored the catabolites addition effects on culture improvement. With the aid of Plackett-Burman and Box-Behnken in Design-Expert 8.0.6, the PB and response surface experiment were designed and analyzed from the single factor results. On the modified GAM medium, the strain concentration was found increasing markedly by 10 times. The addition of some amino acids, vitamins and inorganic salts has contributions for the strain multiplication, especially L-cysteine, VB6 and NaCl. The addition of 55 mg/L of L-cysteine, 20.5 mg/L VB6 and 55 g/L NaCl into the modified GAM medium increased the biomass by 3.59 times compared to the biomass on only modified GAM medium according to the response surface experiment. Through the newly constructed method, we successfully analyzed the amount of *Christensenella minuta* and obtained a novel medium to increase biomass significantly.

**Importance:** *Christensenella minuta* is a heritable bacterium with controversial physiologies associating with both obesity and potential pathogenicity. Since this bacterium is fastidious to culture, it is hard to well understand its biological feature. We develop a strategy for statistical analysis of this low abundant strain and optimize culture condition to make a significant improvement on its biomass and facilitate the researches about the metabolism and function of this bacterium. This work combined the prediction tools and experiments to improve the medium components of *C. munita* and successfully enhance the culturing and increase biomass by more than 10-fold. From this perspective, the project throws some new ideas and also enables access to new knowledge and information in uncultured microbial resources.

## INTRODUCTION

Microbes are the most life forms on the earth and have been applied widely in medicine, environmental protection and other fields. In fact, most microbial lineages have not been isolated in pure culture and it was estimated that only 1% of microbes can be cultured on laboratory media (1). Furthermore, it is still unknown that the abundances and viability of uncultured microbes at different levels of phylogenetic divergence from their cultured relatives. With greater phylogenetic distance correlating with higher levels of evolutionary changes, uncultured groups may have novel undiscovered functions and applications.

Although the culture-independent technologies on uncultured microbes expanding, such as metagenomics and metatranscriptomics, the information about the microbial diversity and distribution can be available. However, the growth requirements and physiological functions of many uncultured microorganisms remain unexplored. For clearly exploring the biological mechanism of individuals from uncultured group, developing the methods on the pure culture, qualitative and quantitative analysis are more realistic (2, 3).

*Christensenella minuta* (*C. minuta*) is a gram-negative gastrointestinal bacterium (4). It was discovered originally associating with obesity through an unknown biological mechanism (5). Most interestingly, the genome of *C. minuta* is highly heritable (6) and presents a valuable application on future obesity therapy (7). Nevertheless, some recent researches have demonstrated that *C. minuta* might be a potential pathogen and have high-risk for its application in the obesity therapy (8). *C. minuta* was isolated from the blood of a patient with a diagnosis of acute appendicitis and this bacterium is one of suspects to cause this disease. From all present studies, the conclusion about the physiological characteristics and pathogenicity of *C. minuta* remain elusive.

*C. minuta* is a strictly anaerobic bacterium (4) and its coefficient of culturing difficulty is extremely high on the laboratory media. Establishing feasible methods for analyzing and culturing *C. minuta in vitro* is conducive to understand its physiological function. We developed a quantifying method using fluorescence quantitative RT-PCR to valuate microbial population; Furthermore, we combined its metabolomic background and positive growth factors to accelerate *C. minuta* growing and increase its biomass. In this paper, our work will be detailed.

## RESULTS

### Specific quantitative measurement of *C. minuta* population basing on fluorescence quantitative PCR

*C. minuta* is hardly to observe when inoculating on the laboratory media and we have trouble to calculate the strain population using conventional quantitative methods such as cell counting, turbidimetry. Quantifying the cellular abundance in our sample is challenging. Since in the microbial culture, there still have some other microbes except *C. minuta*, the specific method for measuring the number of *C. minuta* was our foremost requirement for assessing the bacterial growing improvement.

Fluorescent quantitative real-time PCR (FQ-PCR) has been used widely to quantify the number of genomic copies of microorganisms (9). For distinguishing our target microbe, the method combining specific sequence of microbial 16S rDNA with FQ-PCR is promising to provide a way to quantitate our strain accurately.

From the public databank, 16S rDNA in *C. minuta* is a 1497 bp sequence containing one constant region and one variable region. Its constant region sequence is identified as following: ATCTCAAAAAGCCGGTCCCAGTTCGGATTGTGGGCTGCAACCCGCCCACATGAAGTCGGAGTTGCTAGTAATCGCGAATCAGCATGTCGCGGTGAATGCGTTCCCGGGCCTTGTACACACCGCCCGTCACACCACGGAAGTTGG GAGCACCCGAAGCCAGTGGCTTAACCGTAAGGAGAGAGCTGTCGAAGGTGAGATCAATGACTGGGGTGAAG. After comparing 16S rDNA sequences of three strains from *Christensenella* family, three pairs of primers were designed and sequences were listed in Table 1. To ensure the cloning is specified for *C. minuta*, the electrophoresis and FQ-PCR assessment was performed (Figure 1 and 2). The target band can be observed on *C. minuta* lanes with all primers 1, 2 and 3, but is invisible on *E. coli* lanes and other microbial lanes. As compared to prime 2 and 3, prime 1 had less false positive and smear. From the intensity of cloning and the amplification curve of fluorescence quantification PCR, primer 1 is highly specific and sensitive for cloning 16S rDNA of *C. minuta*.

**Table 1.**
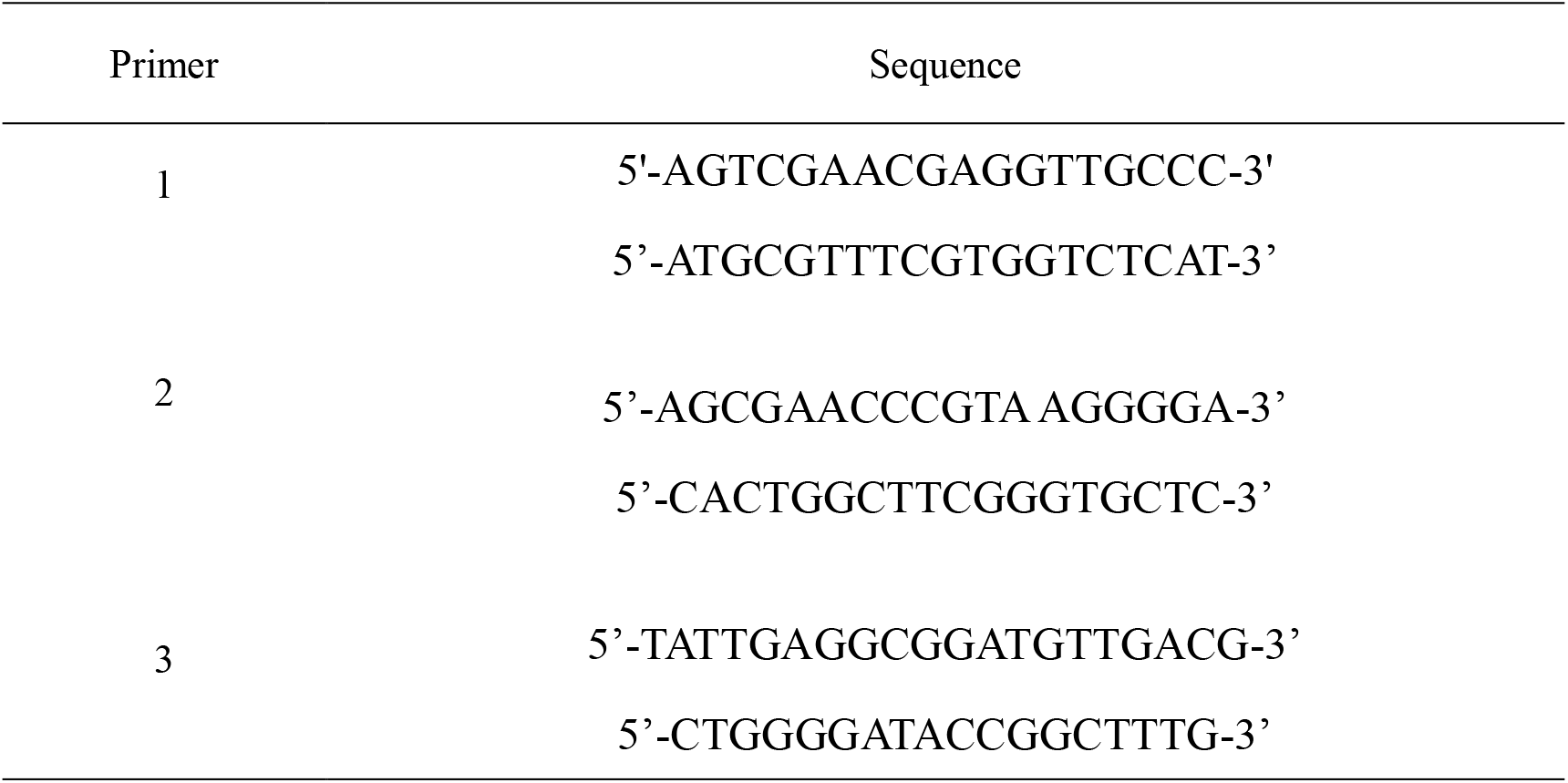
Sequence of specific primers for cloning the constant region in *Christensenella minuta* 16S rDNA

**Fig. 1.**
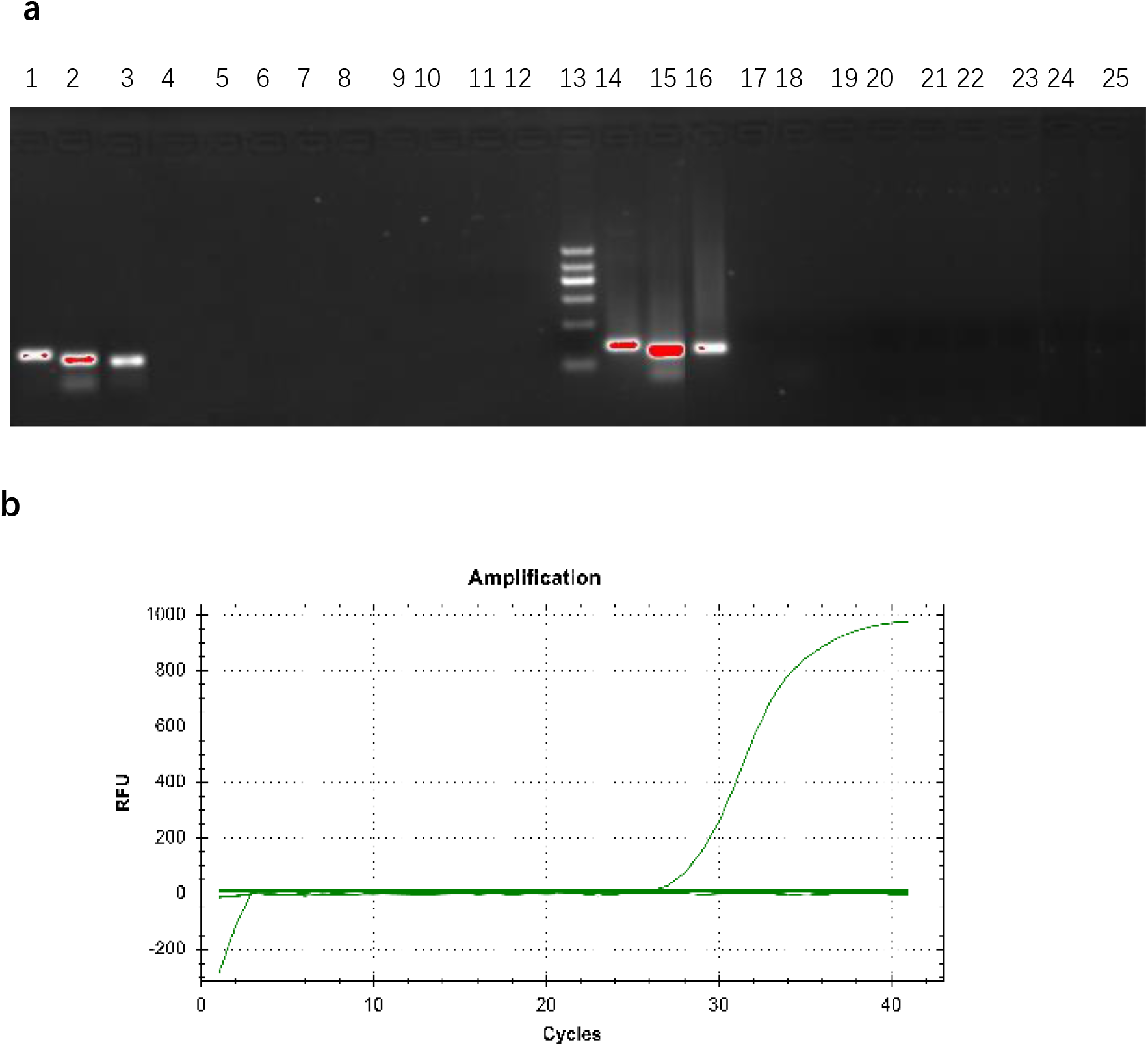
**a** Electrophoretic of PCR products from three pairs of specific primers in different strains. **a** 1~12: Normal PCR reaction; 14~25: Fluorescence quantitative PCR reaction; 13: marker. 1~3 and 14~15: *C. minuta*, primers are 1, 2 and 3; 4~6 and 17~19: *E. coli*, primers are 1, 2 and 3; 7~9 and 20~22: yeast, The primers were 1, 2 and 3; 10-12 and 23-25: soil samples, and the primers were 1, 2 and 3. **b** The amplification of qPCR by primer1. The curve is from *C. minuta*; The straight lines include negative control (ddH20); *E. coli* DH5α; *Agrobacterium* EHA105; *Peptostreptococcus anaerobius*; *Anaerovorax odorimutana*; *Bacillus subtilis*.

**Fig. 2.**
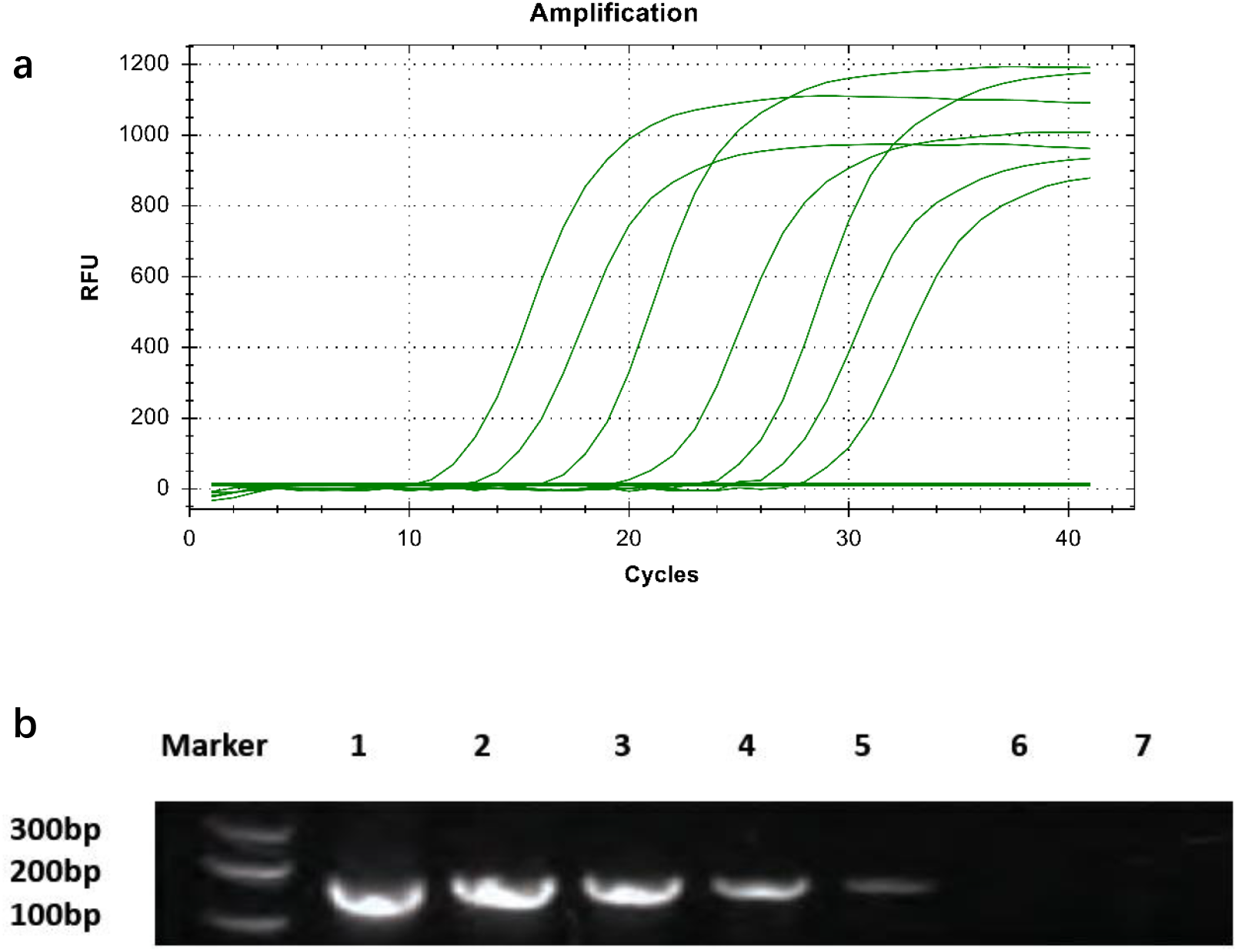
Sensitivity test of FQ-PCR measurement. **a**. The amplification curve of qPCR for *C. minuta* DNA, from left to right: concentration of *C. minuta* DNA is from 5.58×10^10^ copies/μL to 5.58×10^4^ copies/μL. b. Electrophoretic of *C. minuta* DNA, from left to right: DNA marker, C. minuta DNA and concentration from 5.58×10^10^ copies/μL to 5.58×10^4^ copies/μL.

As to the quantitive measurement, the principle is the linear relationship between the *C*_t_ value of each template and the logarithm of the initial copy number of the template is as following.

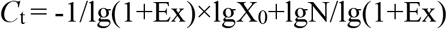

(X_0_ is the initial template amount; Ex is the amplification efficiency; and N is the amount of amplified product when the fluorescence amplification signal reaches the threshold intensity).

In this equation, the initial copy number increases with *C*_t_ value getting lower. A standard curve can be made using a standard with a known initial copy number, where the abscissa represents the logarithm of the initial copy number and the ordinate represents the *C*_t_ value. Therefore, as long as *C*_t_ value of the unknown sample is obtained, the initial copy number of the sample can be calculated from the standard curve. The plasmid cloned by primer 1 from *C. minuta* was extracted and concentration was measured, then the plasmid was cloned on fluorescence quantitative PCR. The calibration curve is constructed by the plasmid concentration logarithm against C_*t*_ value (Figure 3) and will be used in subsequent studies.

**Fig. 3.**
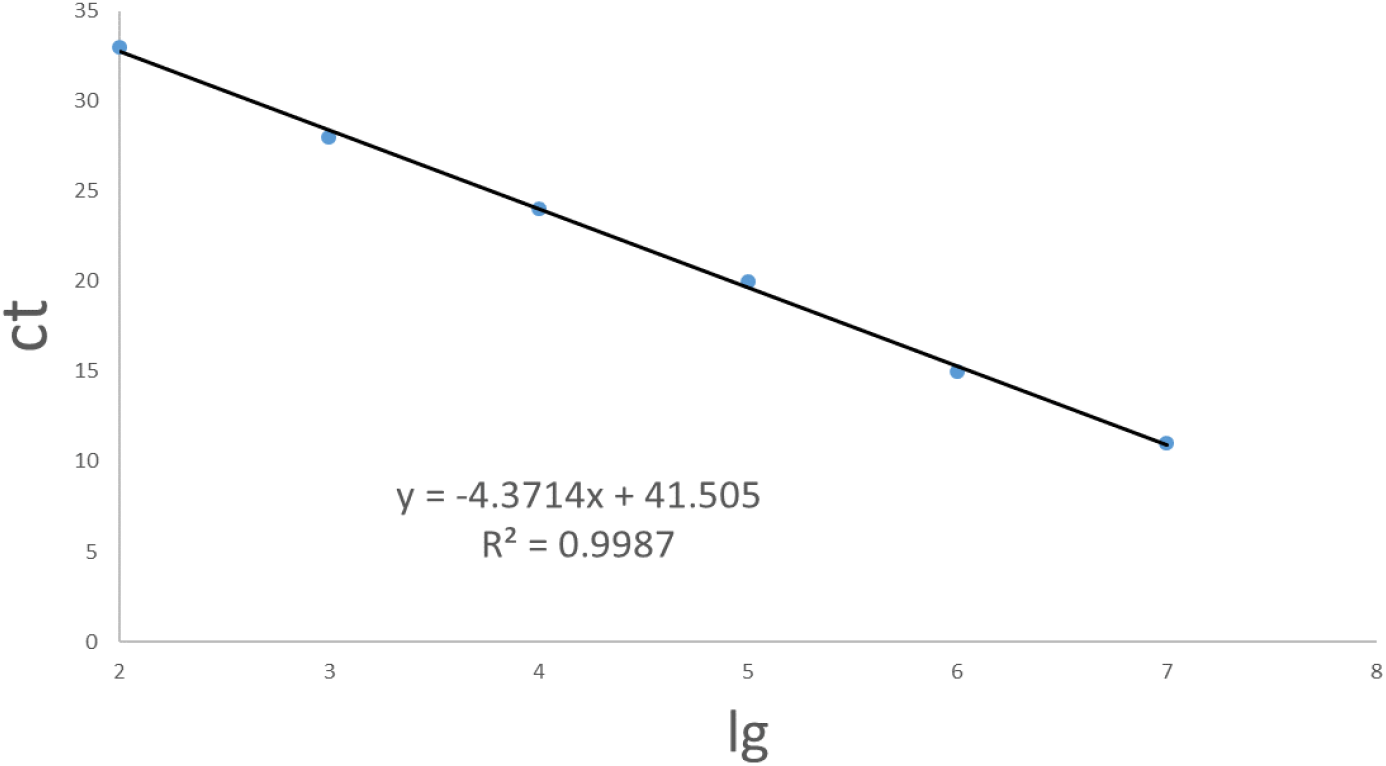
Calibration curve of plasmid concentration logarithm against C*t* value

### Predicting new culturing media with KOMODO

Culturing microorganisms is a challenge that is critical for tapping the biotechnological potential of microbes. For normal microbes, culturing new organisms can be guided by protocols such as Bergey’s Manual of Systematic Bacteriology. However, even with these guides, the culturing for uncultured microorganisms still requires a great deal of trials. A most accepted reason for their unculturability is the absence of key growth factors in artificial media (10), therefore it has a lot of space to improve the media components for culturing fastidious microbes.

The Known Media Database (KOMODO) is a database of microbial media with almost all DSMZ collection (11). KOMODO includes 3,335 media variants, 1,324 metabolic components composing the media and 20,824 media–strain pairings. It has been an online tool that can predict optimized media that microorganisms can grow on by inputing microbial 16S rDNA sequence or an NCBI taxon ID. With the prediction of KOMODO, the options "non-strict aerobic", "inorganic salt in the medium" and "evolution distance is less than 0.04" were defined and 16S rDNA sequence of *C. minuta* (sequence similarity ˃ 85%) was introduced to predict media formula that has improvement on bacterial growing. According to the KOMODO score, three media were recommended including caldicoprobacter medium (score 400), modified GAM medium (score 62.9) and EG medium (score 35.6). *C. minuta* was inoculated on three recommended media to confirm the growth-promoting effect experimentally. The results in Figure 4 have shown that modified GAM medium is the optimal medium for *C. minuta* and the composition of the medium is detailed in Table 2. On the modified GAM medium, the cell amount of *C. minuta* dramastically increased by 16-fold compared to that on basic medium.

**Table 2.**
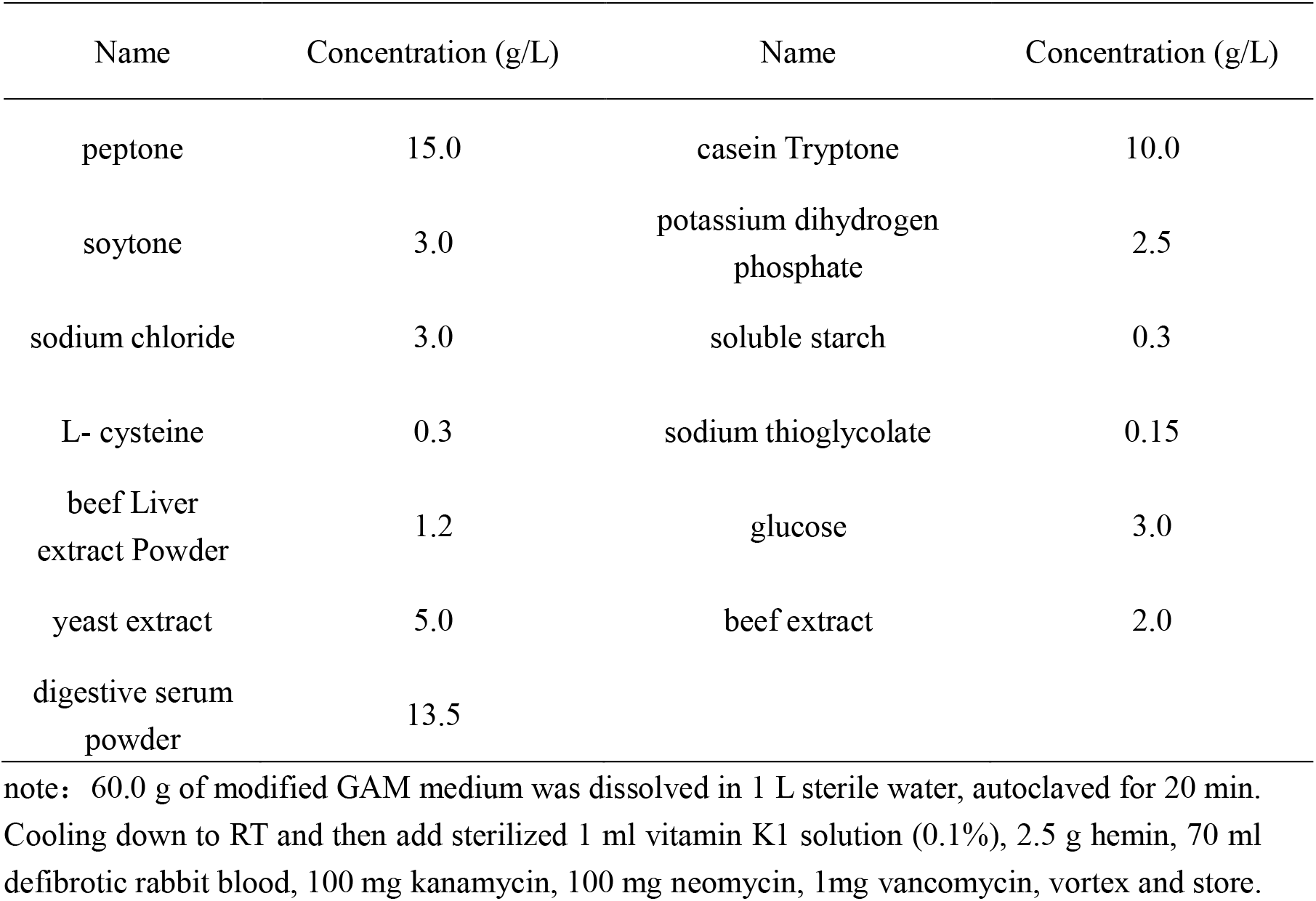
The composition of modified GAM medium (pH 7.2)

| Name | Concentration (g/L) | Name | Concentration (g/L) |
| --- | --- | --- | --- |
| peptone | 15.0 | casein Tryptone | 10.0 |
| soytone | 3.0 | potassium dihydrogen phosphate | 2.5 |
| sodium chloride | 3.0 | soluble starch | 0.3 |
| L- cysteine | 0.3 | sodium thioglycolate | 0.15 |
| beef Liver extract Powder | 1.2 | glucose | 3.0 |
| yeast extract | 5.0 | beef extract | 2.0 |
| digestive serum powder | 13.5 |  |  |

note: 60.0 g of modified GAM medium was dissolved in 1 L sterile water, autoclaved for 20 min. Cooling down to RT and then add sterilized 1 ml vitamin K1 solution (0.1%), 2.5 g hemin, 70 ml defibrotic rabbit blood, 100 mg kanamycin, 100 mg neomycin, 1mg vancomycin, vortex and store.

**Fig. 4.**
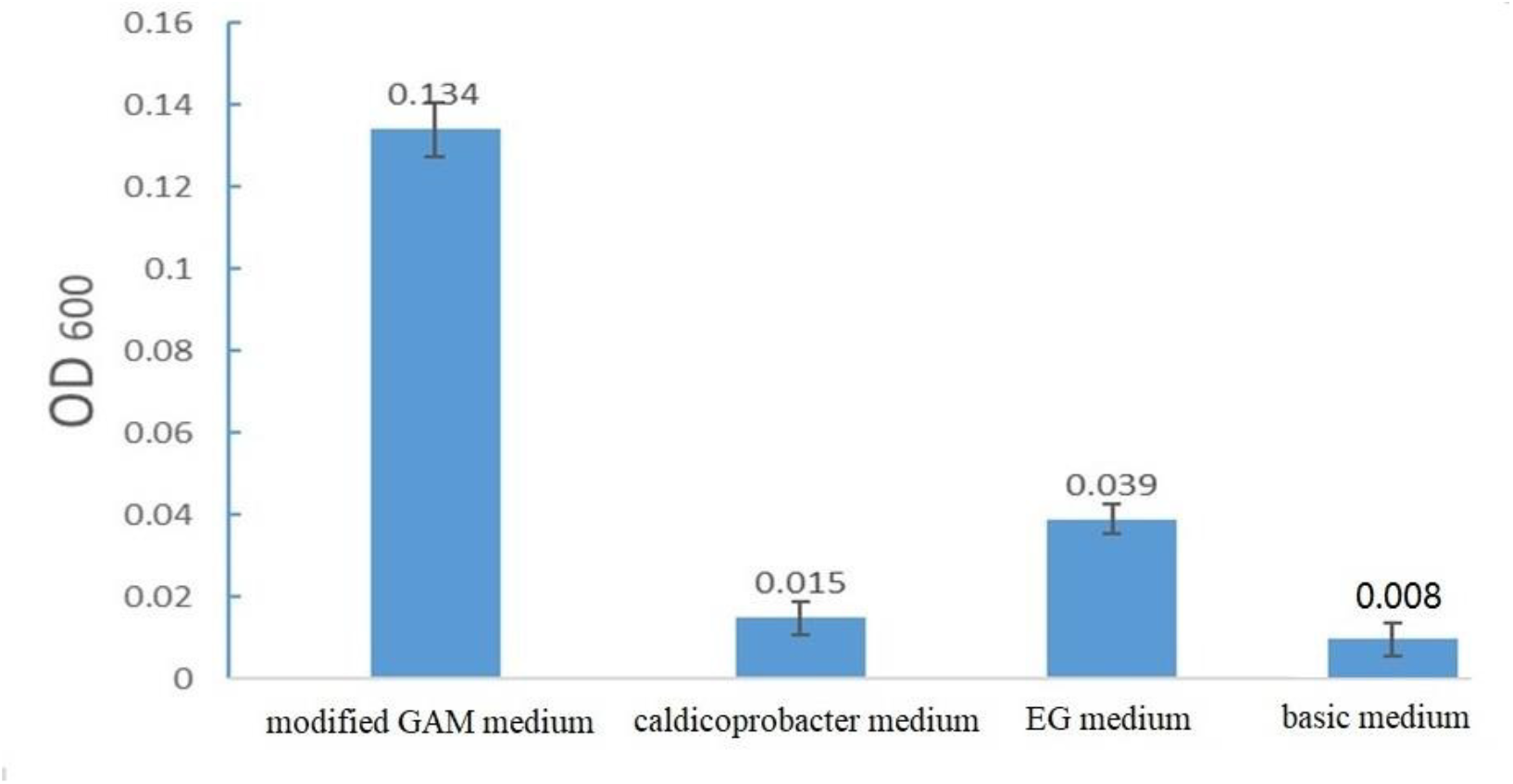
The biomass of *C. minuta* growing on three KOMODO recommended media.

### Medium optimizing basing on metabolomics analysis

In recent years, some culturing efforts, particularly for difficult-to-culture organisms, have begun to include genome and pathway analysis (12,13), as well as high-throughput technologies for determining microbial nutrient needs (14).

KEGG (Kyoto Encyclopedia of Genes and Genomes) is a collection of databases by annotating the reads to known functional gene (15). KEGG is utilized for bioinformatics research including data analysis in genomics, metagenomics, metabolomics and other omics studies. This database provides a comprehensive understanding of the community structure at a high resolution and potential metabolism pathway associated with microbial community.

After the genome sequencing of *C. minuta* was completed, the KEGG map of the carbon metabolism pathway was analyzed and some characteristics *C. minuta* metabolism have been found. Figure 5a shows that in furfural metabolism, the direct pathway of xanthine to amino acids (such as glycine and serine) was substituted by a more complicated path, which may result to the difficult growth of *C. minuta*. Supplementing the medium with the appropriate amount of glycine and serine may increase the culture abundance of *C. minuta*. In the pyrimidine metabolism of figure 5b, it can be seen that UDP, DTP and CTP are the starting materials for the reaction, but the key enzyme (3.6.1.8 ATP pyrophosphatase) lacks. Figure 5c showed the cysteine metabolism of *C. minuta*. In these pathways, the lack of enzymes that converts cysteine and cystine reminded us of supplying cystine to enhance *C. minuta* biomass in culture. Figure 5d shows that the lack of enzymes for the conversion of imidazole-4-acetic acid to aspartic acid in the histone metabolism may also be responsible for the low abundance of culture of *C. minuta*. The product aspartic acid may increase the culture abundance of *C. minuta*. In Figure 5e, the amino acid and nucleotide sugar metabolic pathways of *C. minuta* lack the enzyme (UDP-glucuronate 5’-epimerase) for the conversion of UDP-GleA (glucuronide) to UDP-L-IdoA. Figure 5f shows that in the metabolism of *C. minuta*, VB12 is an important intermediate in the reaction. There are two synthetic pathways, but none of the related enzymes 1.16.1.6 (cyanocobalamin reductase) or 1.16.1.4 (cob(II) alamin reductase). If VB12 is added directly to the culture medium, it is expected to improve the culturability of *C. minuta*. The predicted results in Figure 5g show that *C. minuta* lacks an important pathway in the VB6 metabolism of pyridoxine, isopyridoxal, hydroxymethyl succinate semialdehyde, which lacks key enzymes 99.9 (pyridoxine 5-dehydrogenase), 1.14.12.5 (5-pyridoxate dioxygenase), 3.5.1.66 (2-(hydroxymethyl) −3-(acetamidomethylene) succinate hydrolase), the deletion of this pathway gene may also lead to the difficulty of *C. minuta* in culturing, if the key metabolites in the above pathway are added to the medium, the culture abundance may be increased.

**Fig. 5.**
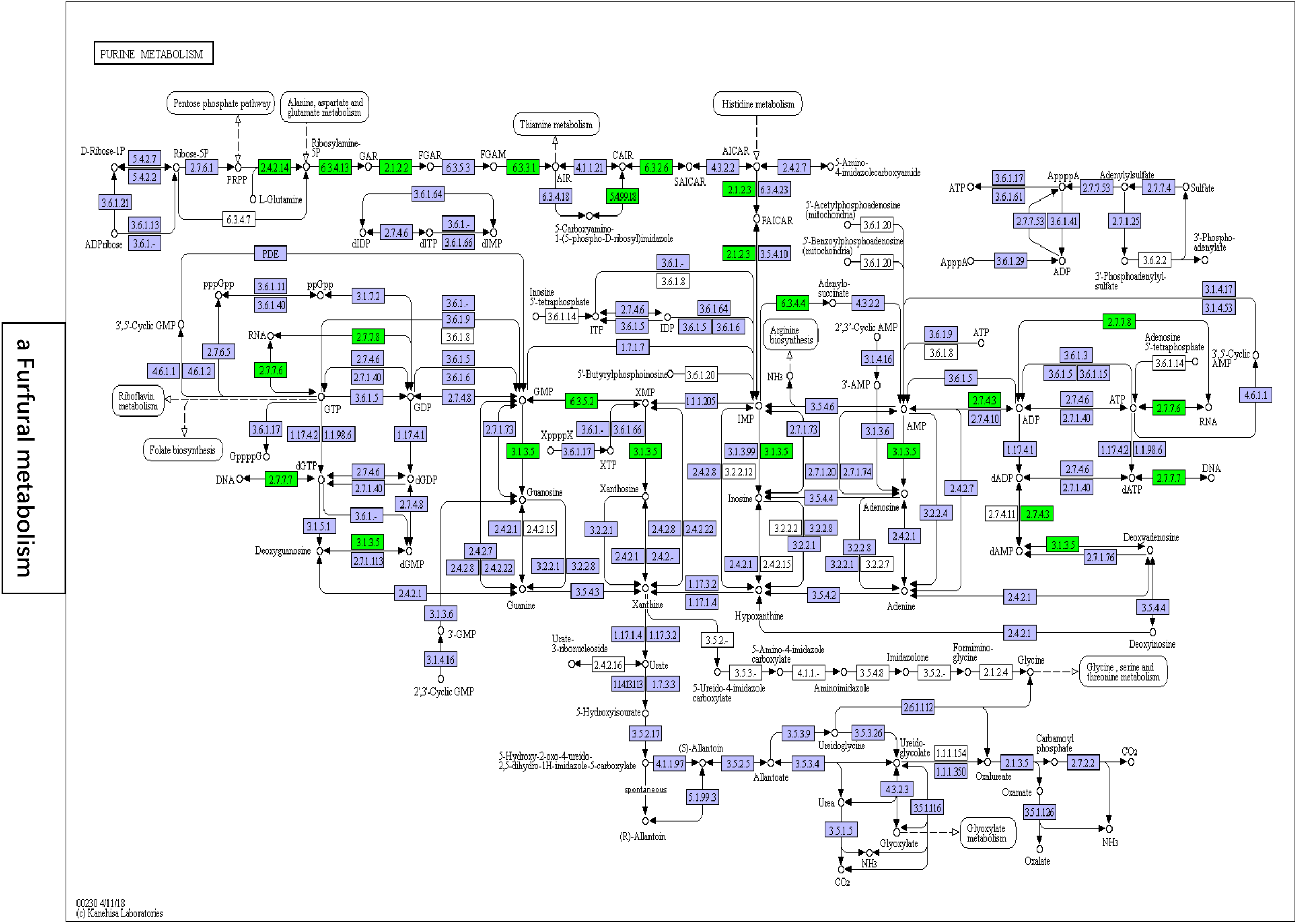
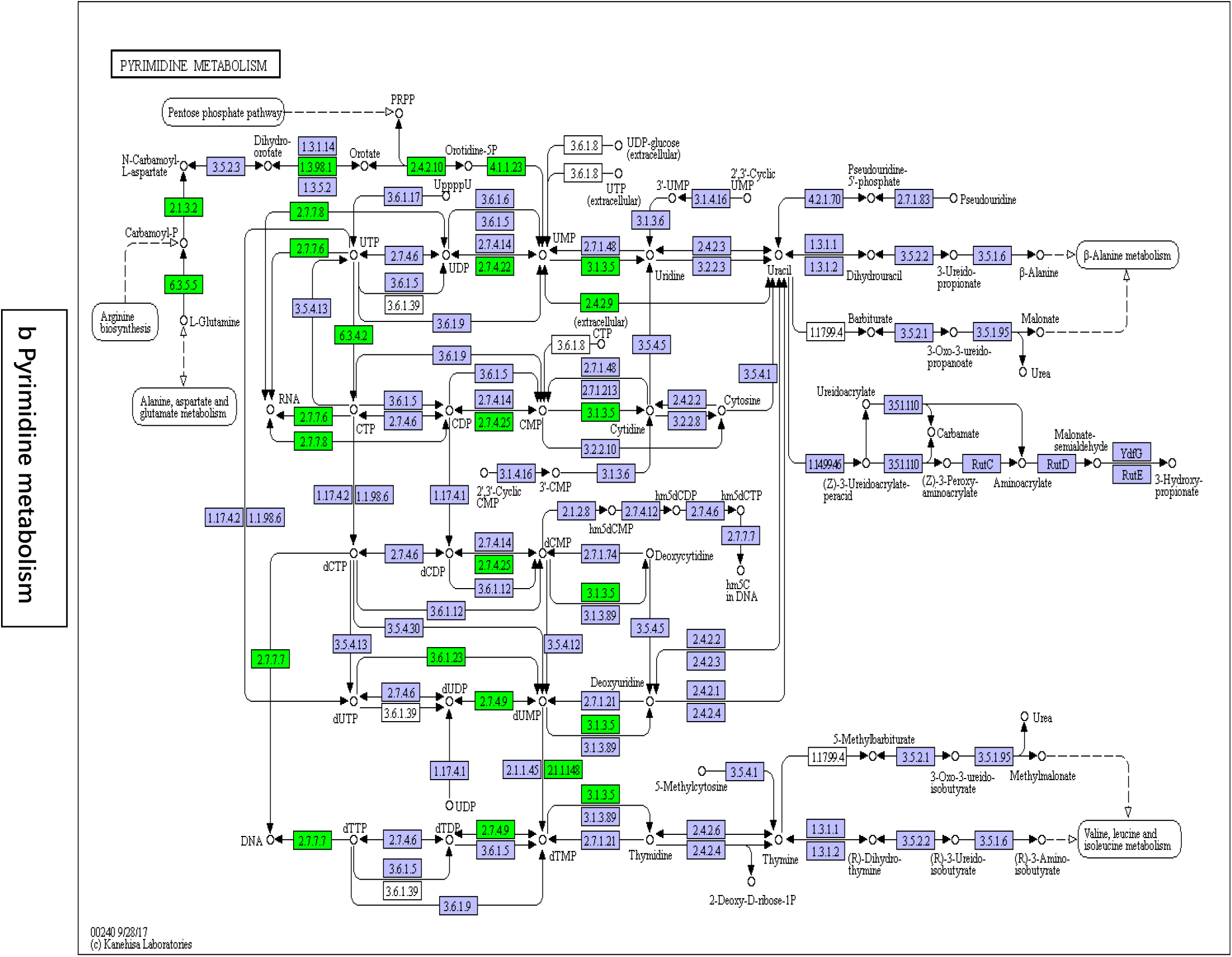
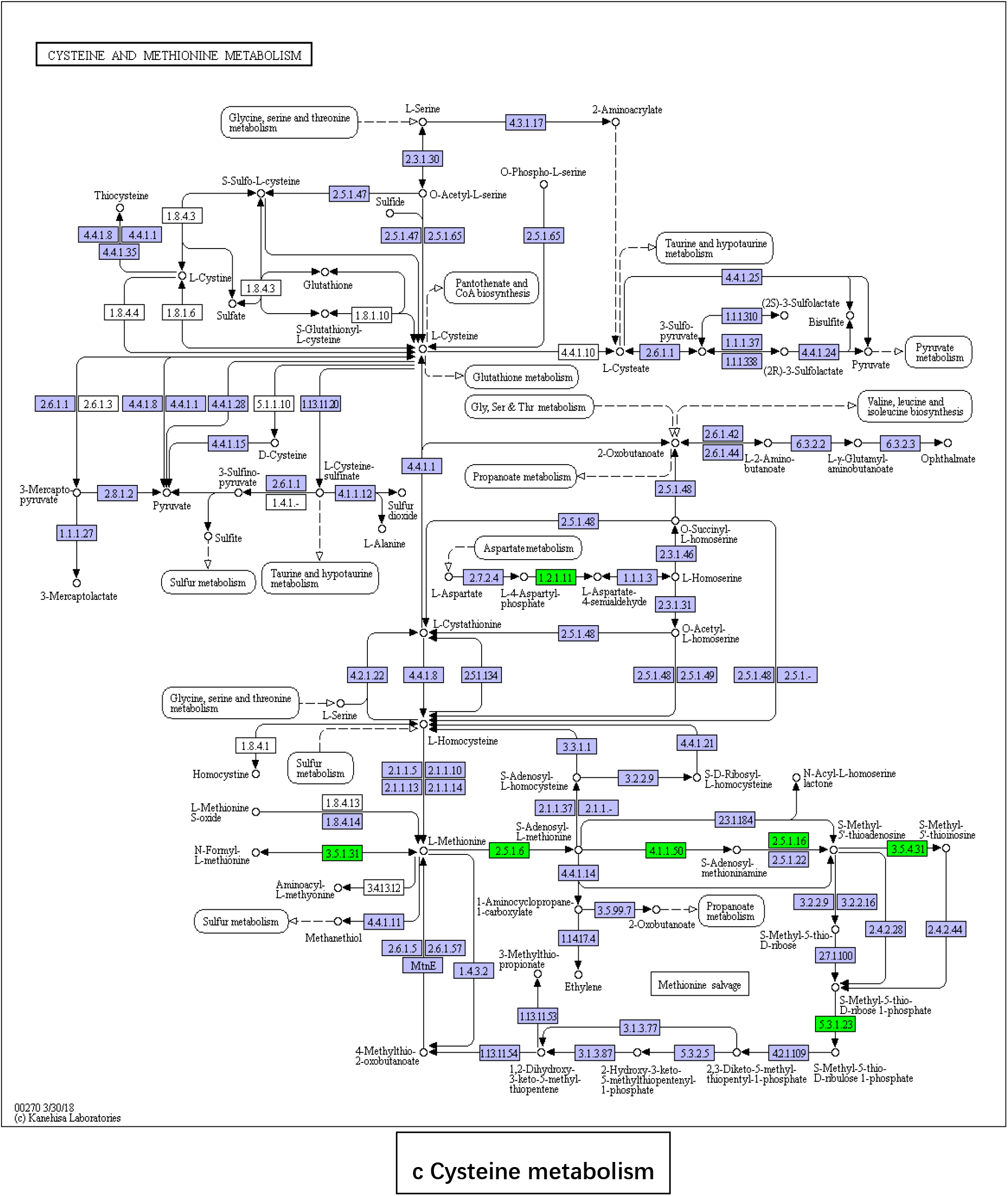
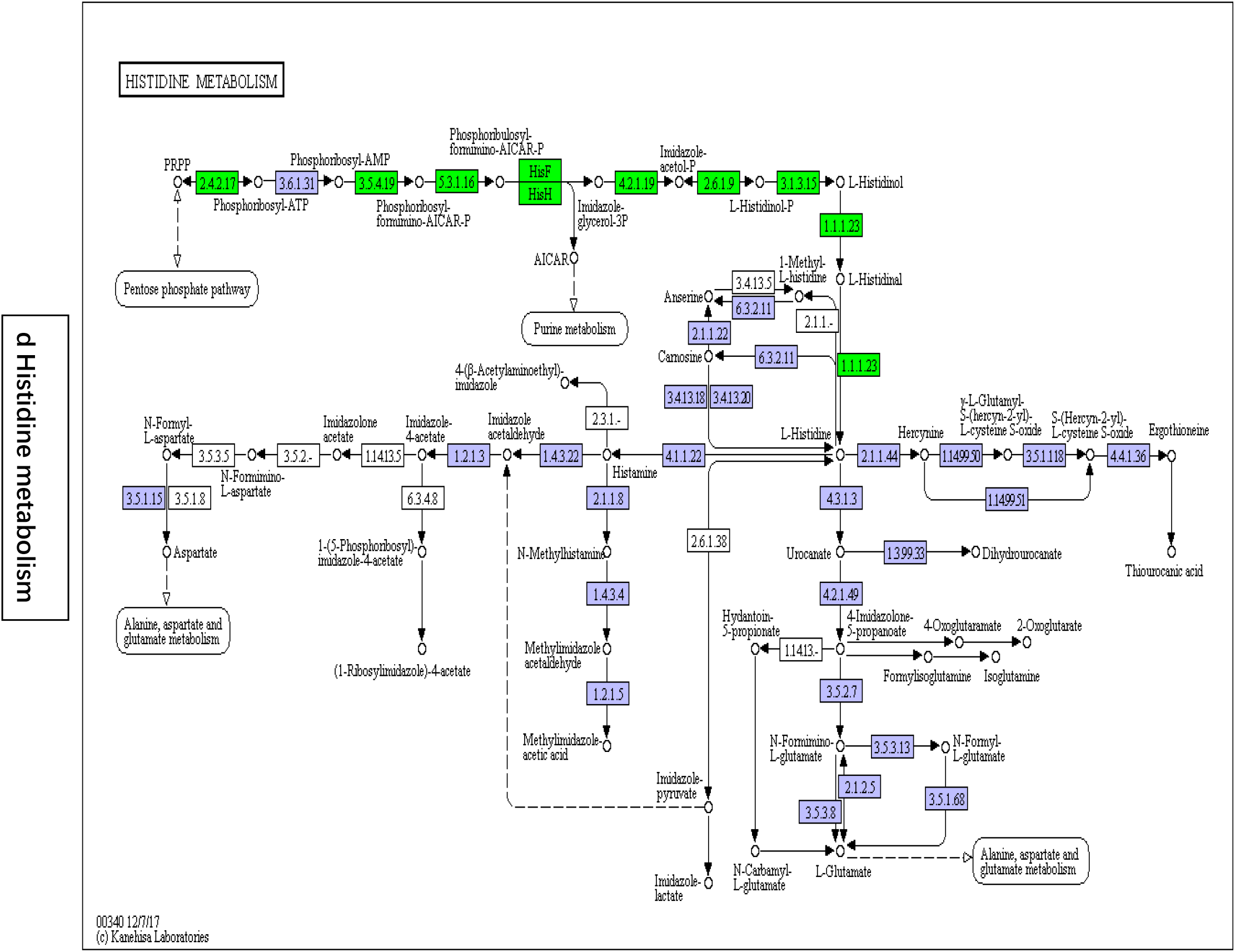
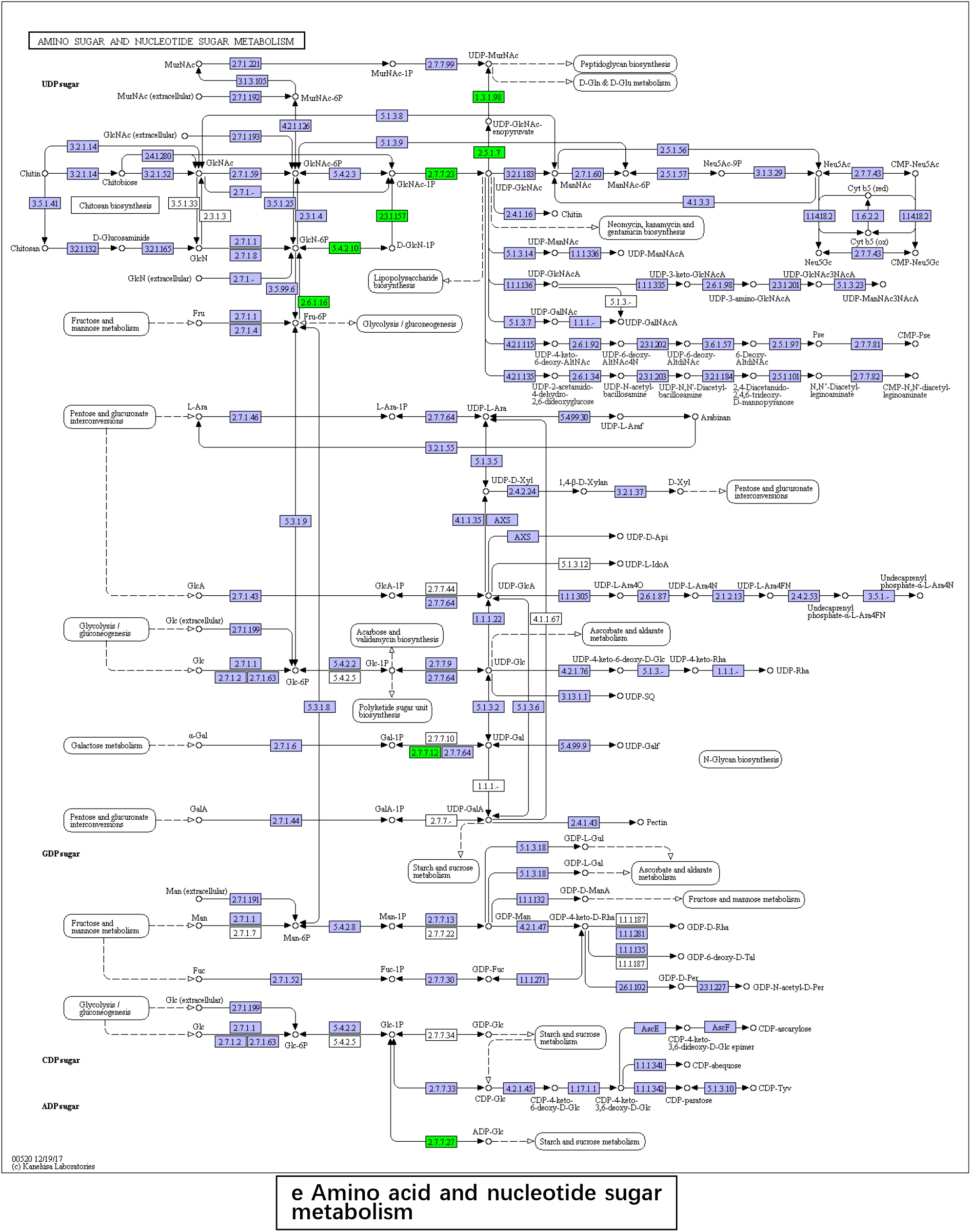
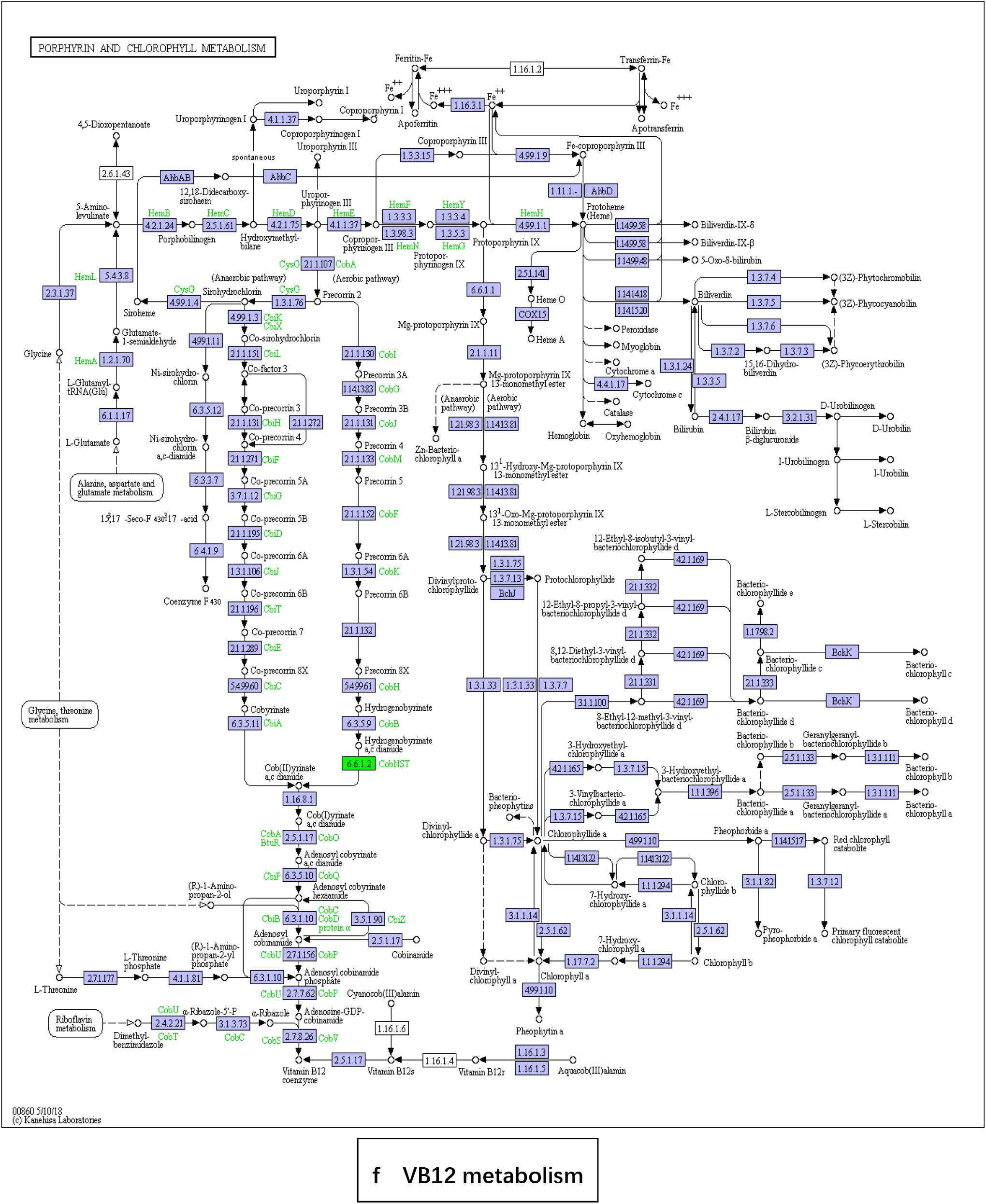
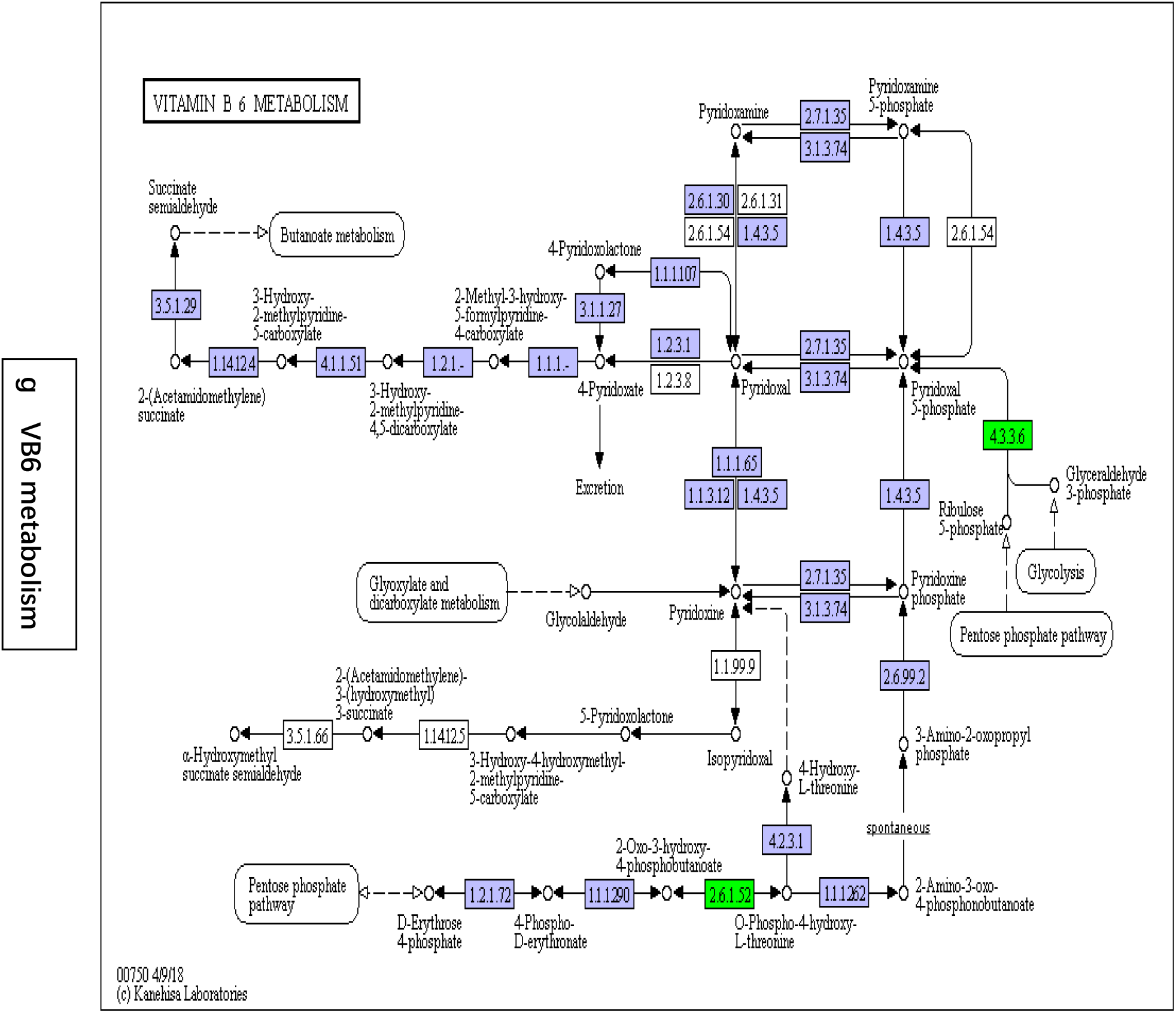
Metabolome map of *Christensenella minuta*

According to the metabolic pathway map analyzed above, serine and glycine were selected to verify the metabolism of furfural, cystine and cysteine to verify cysteine metabolism, aspartic acid to verify histidine metabolism, UMP, CMP to verify pyrimidine Metabolism, VB12 verifies VB12 metabolism, and VB6 verifies VB6 metabolism. Refer to the amino acid concentration in DMEM medium (L-serine 42.00 mg / L, glycine 30.00 mg / L, L-cystine 63.00 mg / L, L-HCl cysteine 80.00 mg / L, L-winter The amino acid was added at 16.00 mg/L, and the VB6 (pyridoxine hydrochloride) in this formulation was 4.00 mg/L. Vitamins were added with reference to human serum VB12 concentration (70 to 590 pmol/L) and VB6 (14.6 to 72.9 nmol/L). UMP and CMP were added with reference to yeast synthesis UMP concentration (14.3 g/L). As shown in Table 3, after adding each test substance to the medium, the growth of *C. minuta* showed a certain promotion effect (Multiples > 1).

**Table 3.** Effects of each test substance on the proliferation of *Christensenella minuta*

| Test substance | Conc (mg/L) | Multiples | Conc (mg/L) | Multiples | Conc (mg/L) | Multiples |
| --- | --- | --- | --- | --- | --- | --- |
| L-serine | 100 | 1.75 | 40 | 1.65 | 10 | 1.79 |
| L-glycine | 100 | 1.29 | 30 | 1.75 | 10 | 1.78 |
| L-cystine | 100 | 1.79 | 60 | 1.45 | 10 | 1.11 |
| L-cystine | 100 | 1.03 | 80 | 1.87 | 10 | 1.64 |
| L-aspartate | 100 | 1.67 | 20 | 1.52 | 10 | 1.34 |
| 5-uridine monophosphate disodium salt (UMP) | - |  | 10000 | 1.35 | 1000 | 1.24 |
| Cytidine-5-disodium phosphate (CMP) | - |  | 10000 | 1.67 | 1000 | 1.56 |
| Pyridoxine hydrochloride (VB6) | 40 | 0.87 | 4 | 1.73 | 1 | 1.66 |
| Cobalamin (VB12) | 40 | 0.46 | 4 | 1.38 | 1 | 1.59 |

Once the basic culture conditions were determined by single factor experiments, PB experiments were performed to determine which factor concentration changes had a significant effect on the multiples of *C. minuta*. The results are shown in Table 4 that the three most important factors among the many influencing factors are L-Cysteine, VB6 and NaCl, the corresponding P-value values were 0.0001, 0.0006 and 0.0036, respectively, indicating a significant difference.

**Table 4.** Analysis of variance of selected factors

| Source | Sum of Squares | df | F Value | p-value | Prob > F |
| --- | --- | --- | --- | --- | --- |
| L-serine | 3.234E-003 | 1 | 19.91 | 0.0467 |  |
| L-glycine | 4.294E-003 | 1 | 26.44 | 0.0358 |  |
| L-cystine | 4.840E-003 | 1 | 29.80 | 0.0320 |  |
| L-cystine | 0.28 | 1 | 1738.98 | 0.0006 | significant |
| L-aspartate | 3.710E-003 | 1 | 22.84 | 0.0411 |  |
| UMP | 0.023 | 1 | 139.27 | 0.0071 |  |
| CMP | 0.014 | 1 | 88.37 | 0.0111 |  |
| VB6 | 1.20 | 1 | 7366.10 | 0.0001 | significant |
| L-NaCl | 0.045 | 1 | 277.18 | 0.0036 | significant |

The response surface test was carried out with concentrations and combinations of three factors with significant differences of NaCl, L-cysteine and VB6. The results (Table 5) showed that the addition of 55 mg/L of L-cysteine, 20.5 mg/L VB6 and 55 g/L NaCl into the modified GAM medium were optimized and improved the growth of *C. minuta*, which increased the biomass of strains by 3.59 times, with significant differences.

**Table 5.** Response surface experiment result

| Run | Factor 1<br>L-cystine<br>mg/L | Factor 2<br>VB6<br>mg/L | Factor 3<br>NaCl<br>g/L | Multiples |
| --- | --- | --- | --- | --- |
| 1 | 10.00 | 20.50 | 10.00 | 3.15 |
| 2 | 55.00 | 20.50 | 55.00 | 3.53 |
| 3 | 55.00 | 20.50 | 55.00 | 3.54 |
| 4 | 100.00 | 20.50 | 100.00 | 3.38 |
| 5 | 10.00 | 40.00 | 100.00 | 3.21 |
| 6 | 55.00 | 1.00 | 10.00 | 3.23 |
| 7 | 100.00 | 20.50 | 10.00 | 3.42 |
| 8 | 55.00 | 20.50 | 55.00 | 3.59 |
| 9 | 10.00 | 20.50 | 100.00 | 3.25 |
| 10 | 55.00 | 1.00 | 100.00 | 3.18 |
| 11 | 100.00 | 40.00 | 55.00 | 3.39 |
| 12 | 55.00 | 20.50 | 55.00 | 3.56 |
| 13 | 100.00 | 1.00 | 55.00 | 3.3 |
| 14 | 55.00 | 40.00 | 100.00 | 3.42 |
| 15 | 10.00 | 1.00 | 55.00 | 3.08 |
| 16 | 55.00 | 40.00 | 10.00 | 3.43 |
| 17 | 55.00 | 20.50 | 55.00 | 3.57 |

The significance of the above factors is analyzed by the contour and 3D graphs of the AB, AC, and BC interaction results. The concentric center of the contour (the red dot in the graph) falls within the common values of the factors AB, AC, and BC (Fig. 6). Therefore, the maximum value of the dependent variable was obtained in the range of the common concentration, indicating that the optimal composition of the *C. minuta* medium was optimized in this experiment.

**Fig. 6.**
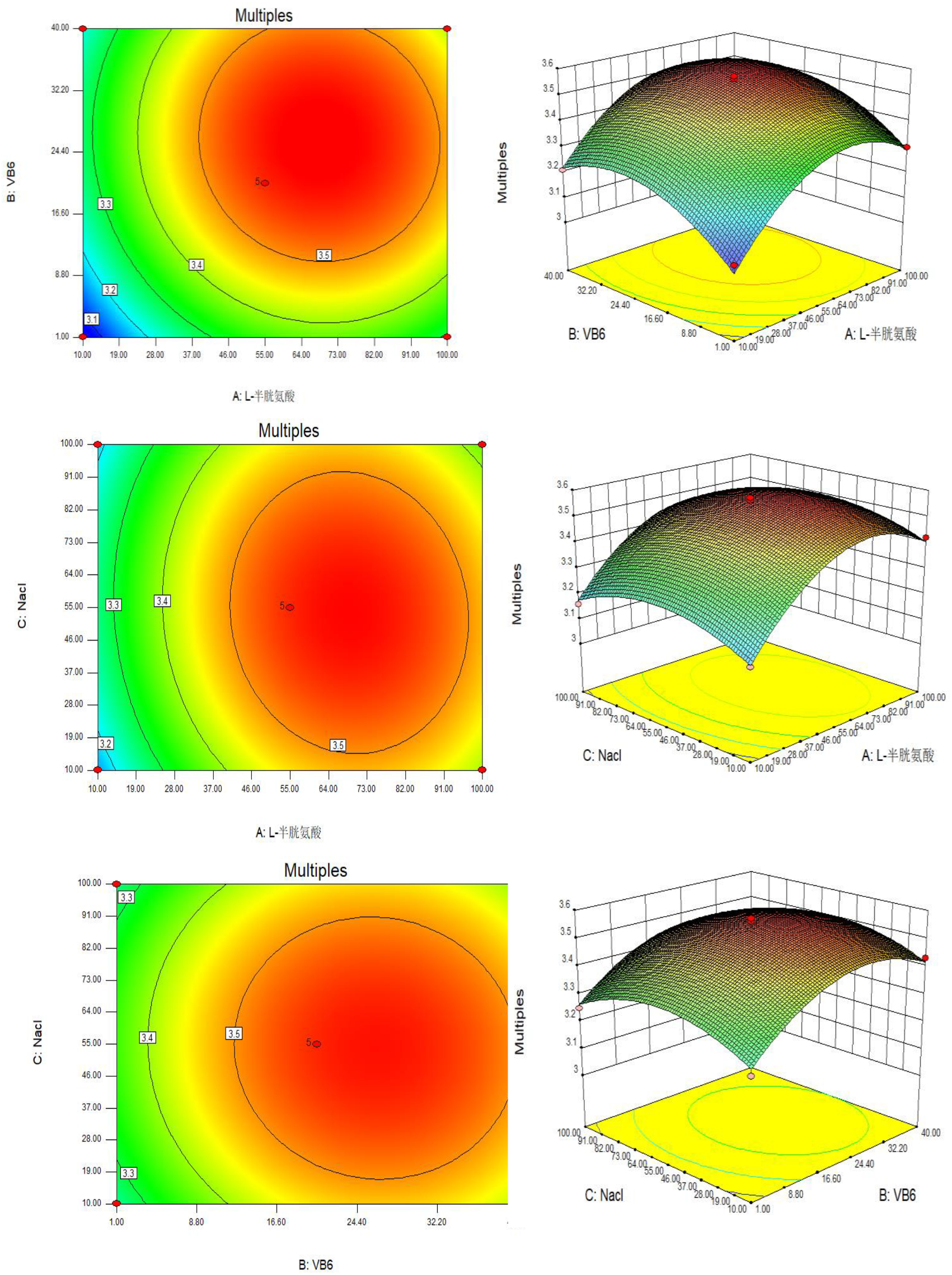
Contour and 3D plots of significant results for AB, AC, and BC interactions

## CONCLUSION

At present, the microorganisms that can cultivate in the laboratory account for only 1% of the microbial species, which means most of the microorganisms in nature cannot be cultured in the laboratory. Since that, most microbes cannot be scientifically studied. Although the development of metagenomics, transcriptomes, etc., has enabled researchers to conduct taxonomic and gene-based informatics analysis of difficult-to-cultivate or non-cultivable microorganisms, it is still necessary to conduct functional research and resource development for a specific microbial individual. In the development of microbial culture methodology, scientists can only conduct in-depth research on target microbial resources on the basis of cultivable conditions.

In this work, we combined the prediction tools and experiments to improve the media components of *C. munita* and successfully enhance the culturing and increase biomass by more than 10-fold. From this perspective, the project throws some new ideas and also enables access to new knowledge and information in uncultured microbial resources.

In addition, microbes are the source of important biosynthetic resources. Uncultivated microorganisms not only contain a large number of unexploited material resources in terms of quantity and species, but these microbial populations that are not recognized by humans are a treasure house. Providing new functional molecules for biological applications, and has great potential for development in new drugs and new enzymes.

## MATERIALS AND METHODS

### *Christensenella minuta* and its basic culture

*C. minuta* was obtained from feces sample. The single colony was screened and identified with 16S rRNA. The strain was basically cultured on Gifu anaerobic medium (Nissui Pharmaceutical). Each inoculated sample was incubated at 37°C for 3 days in an anaerobic glovebox (Coy Laboratory Products), which contained 88% nitrogen, 7% hydrogen and 5% carbon dioxide.

### Cloning of specific fragments in *C. minuta* and PCR product transforming

The gene of *C. minuta* was extracted with TIANGEN MiniElute DNA Kit (DP316) and 16S rDNA fragment was cloned with the template of total DNA by specific designed primers. PCR procedure was followed as the introduction in QIANGEN PCR kit (VI102). PCR reaction is 95°C 5 min; 95°C 30 s, 60°C 30 s, 72°C 30 s, 30 cycles; 72°C 10 min. pClone007 Blunt Simple Vector Kit was used for the gene linking to make recombinant plasmids. The recombinants were transformed into *E. coli* competent and the procedure was shown in Figure 7.

**Fig. 7.**
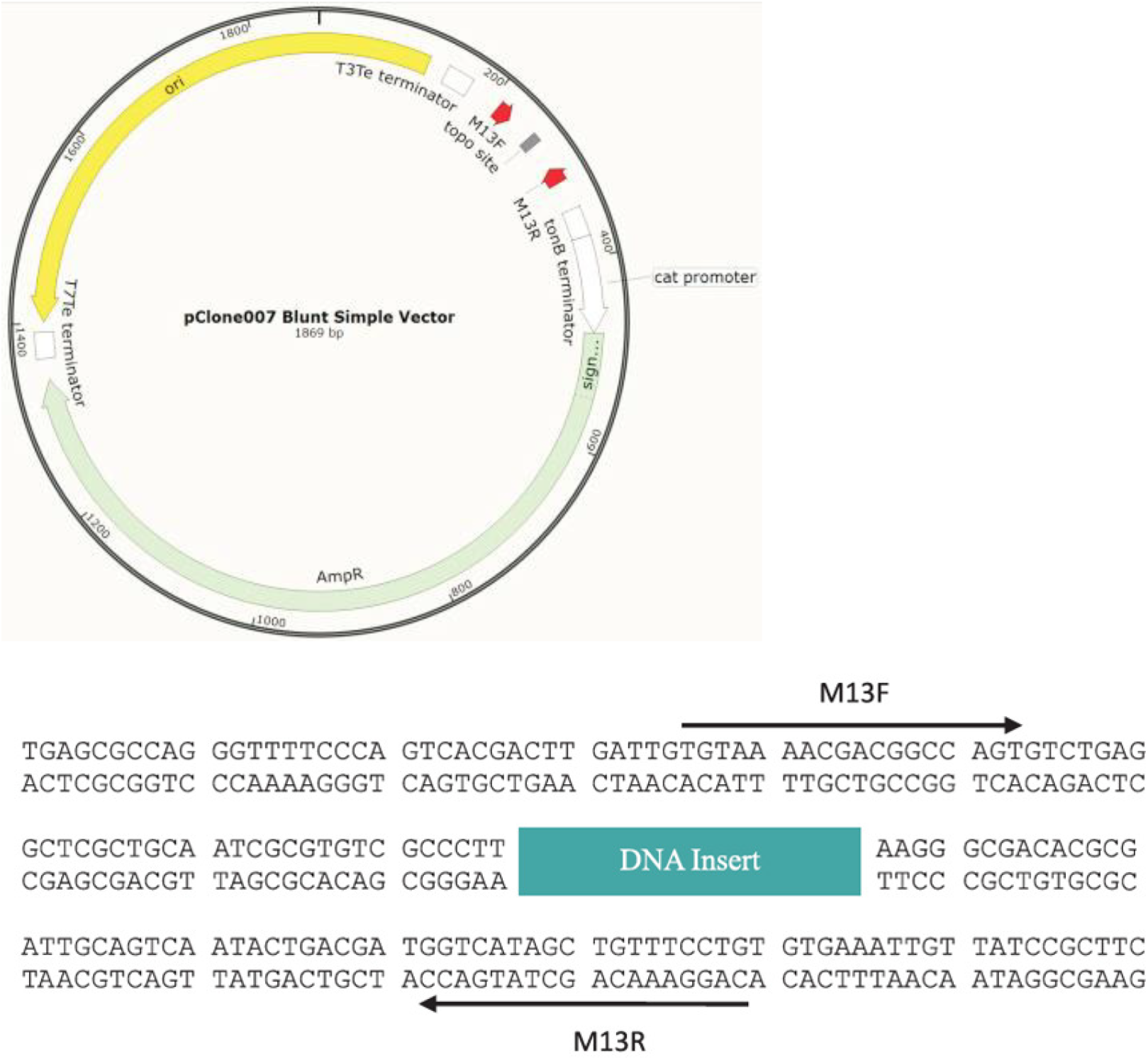
The map of pClone007 Blunt Simple Vector and its link with 16S rDNA of *C. minuta*.

### Fluorescence quantitative PCR and calibration curve construction

All recombinants were extracted and concentration was measured. A 10-fold gradient dilution of the positive plasmid was used to construct a calibration curve against *C*_t_ value from the fluorescence quantitative PCR. Use 2×T5 Fast qPCR Mix(SYBRGreenI) to perform qPCR and the reaction system configuration is 2×T5 Fast qPCR Mix 10 μL, upstream and downstream primer 0.4 μL respectiveyly, 50×ROX reference dye 0.4 μL, DNA sample 2.0 μL and 6.8 sterile water. PCR reaction procedure is 95°C 30 s; 95°C 15 s, 61°C 60 s, 40 cycles. Fluorescence was detected in each cycle.

### KOMODO prediction and metabolite analysis

On Komodo page (http://delta-tomcat-vm.cs.tau.ac.il:40678/Komodo/growrec.htm), the optimal media were predicted for *C. minuta*. The conditions were limited: Is organism Aerobic (Yes/No/Unknown): No; Does Organism grow in Saltly Media (Yes/No/Unknown): Yes; Maximal phylogenetic distance (range:0.0 – 1.0, default:0.04): default。16S rRNA data of *C. minuta* was inputted and blast ‘Identities’ Low Limit %: 85%. *C. minuta* was inoculated on prediceted media from KOMODO database. After culturing for 7 days, the samples were determined by fqPCR to evaluate the optimal medium.

The metabolite analysis was performed with KEGG (https://david.ncifcrf.gov/home.jsp;) combining genomics data of *C. minuta* from NCBI (https://www.ncbi.nlm.nih.gov/). Choose “GO ONTOLOGY and PATHWAY” and the prediction results were obtained. *C. minuta* was inoculated on modified GAM media supplying with metabolites predicted from KEGG. After culturing for 7 days, the samples were determined by fqPCR to evaluate the optimal medium.

## Acknowledgements

This research was supported financially by Hunan Provincial Natural Science Foundation of China (2016JJ4038) and Double first-class construction project of Hunan Agricultural University (SYL201802002).

